# An RNA sequence that reprograms ribosomes to bypass a 50 nucleotide coding gap is encoded by a mobile element whose sequence conservation illuminates its bypass mechanisms

**DOI:** 10.1101/2022.08.31.505936

**Authors:** Gerard Manning, John F Atkins

## Abstract

**Background:** A remarkable sequence in phage T4 causes ribosomes to skip over a 50 nucleotide insert within a topoisomerase subunit gene, and resume correct synthesis of the protein at a high efficiency. Its mechanism has been extensively studied but it remained an isolated phenomenon whose origin and full function are still a mystery.

**Results:** We have found dozens of homologous cases in genomic and metagenomic sequences, all part of a mobile DNA element that repeatedly inserts in topoisomerase genes of Myoviridae phages. These have substantial sequence diversity, with selective conservation that specify the elaborate set of mechanisms found experimentally to underlie this extreme case of translational recoding. These sequences provide new variations on these mechanisms, and introduce additional features that may also be important for bypassing. These include a series of RNA secondary structures, a conserved stop codon or rare ‘hungry’ codon at the start of the bypass, a Shine-Dalgarno sequence flanked by AU-rich sequence, and residues in the nascent peptide that prime the ribosome for bypassing.

**Conclusions:** These data provide an evolutionary foundation for the experimentally derived mechanisms, highlight several new features of the sequence, and provide substantial new variations on the bypass theme that will allow further experimental exploration of biologically meaningful variants.

## Introduction

Protein synthesis is remarkably processive, allowing the accurate synthesis of proteins as long as human titin (>35,000 codons), with minimal loss of reading frame. However, programmed shifts in reading frame are widespread and used to make alternative proteins and fulfill a variety of regulatory functions. These shifts shed light on the mechanism of translation, the annotation of genomic sequences, and in some instances on regulatory features [1–4].

Detachment of a peptidyl-tRNA anticodon from its paired mRNA can be followed by its productive re-pairing within the same ribosome to a different but matched, sequence in the mRNA. When the re-pairing is to a codon that does not overlap the codon from which the pairing dissociated, the term translational bypassing is often used (reviewed in [1–3]). This is in contrast to the much more common occurrence of frameshifting where the reading frame is shifted by +1, −1, or −2.

The first discovered bypassing occurrence [5], and similar ones shortly afterwards [6–8] involved “hopping” over a slow-to-decode codon with 9 nucleotides specifying a single amino acid. As the re-pairing of the tRNA anticodon to mRNA involved is frame independent, these cases were similar in principle to later studied long distance, non-programmed and hence also low efficiency, synthetic instances [9, 10]. In those occurrences, the level of aminoacyl-tRNA for the slow-to-decode codon was seriously limiting.

A similar case involving limitation of an aminoacyl-tRNA was recently reported in human cancer model, where interferon-γ from attacking immune cells caused a depletion of aminoacylated tryptophan tRNA in melanoma cells. This caused ribosomal stalling at tryptophan codons progressing to downstream out-of-frame codons, with consequences for protective immune system display of aberrant peptides [11]. In the initial aberrant peptides characterized, the re-pairing to mRNA was at an overlapping codon and so while bypassing is the sense used here was not involved, further work may yet reveal its occurrence. However, there is suggestive evidence for natural utilization of bypassing, for a regulatory purpose by a *Streptomyces* phage [1, 12].

However, more relevant to the present work is the well-established natural programmed bypassing in mitochondrial decoding by the yeast *Magnusiomyces capitatus* [13–15]. *M. capitatus* contains disruptive inserts, byps, in 12 out its 15 essential mitochondrial protein coding genes, with up to 12 inserts in a single gene. It seems at least highly likely that the translational bypassing that permits the synthesis of functional products is both precise and highly efficient [13, 14]. Such inserts are frequent, there being a total of 81 and they range from 27 to 55 nt in length. The inserts bypassed seem to have spread like ‘GC cluster’ miniature mobile elements, though with bypassing ability and so viability of cells with the inserts being dependent on some unknown features specific to these yeasts [13, 14].

Here we focus on the first identified instance of programmed bypassing [16, 17]. Ironically the gene in whose decoding this major departure from sequential triplet decoding occurs, gene 60 of phage T4, is almost adjacent to the gene employed to reveal successive non-overlapping ‘triplet’ decoding [18]. T4’s ancestral topoisomerase 2 large subunit encoding gene was first split into two, genes *39* and *60*, with *gene 60* containing an additional frameshifting insertion of 50 nt between codons 46 and 47 (Fig 1) [16, 19]. Close to half the ribosomes bypass this 50 nt coding gap insert. These ribosomes pause protein synthesis at the ‘take-off’ codon, Gly46 (GGA), where the peptidyl-tRNA dissociates from its mRNA but is retained within the ribosome. After pseudo-translocation the retained peptidyl-tRNA re-pairs to the mRNA at a matched GGA glycine codon at the end of the coding gap and protein synthesis resumes at the 3’ adjacent codon. This unique behavior has been extensively studied and revealed several signals, in both the RNA and the nascent peptide, working together to enable the bypass (Fig 1) [17, 20–32].

**Figure 1.**
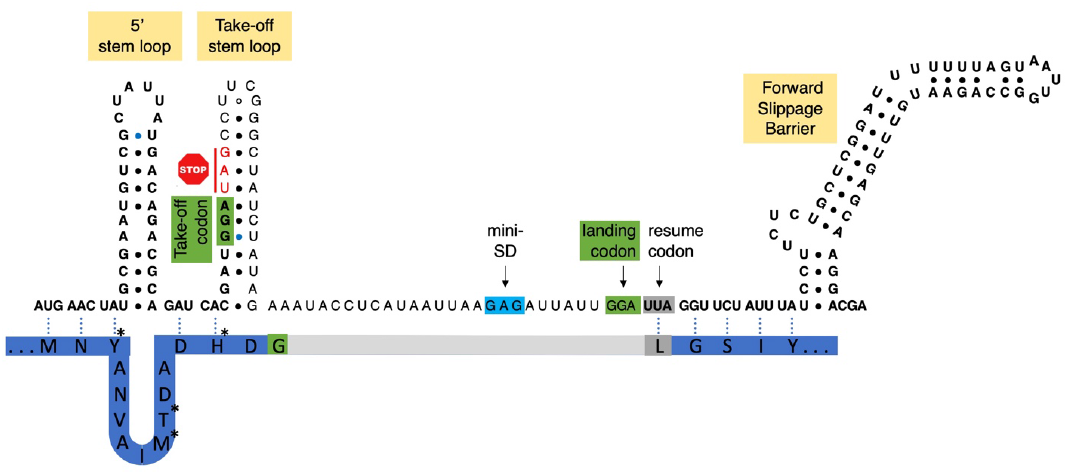
Schematic of the T4 gene 60 bypass region. Coding sequence is in bold, 50 nt bypass cassette is in light font. Translation pauses at the take-off codon (green) which is followed by a stop codon, and resumes after the landing codon (green). This is facilitated by the upstream 5’ and take-off stem loops, and by the downstream Forward Slippage Barrier. Partial ORF2 peptide shown with dashed lines linking codons and corresponding residues, asterisks mark key residues that drive bypassing by twisting the ribosome as it approaches the take-off, grey indicates coding gap and take-off and resume residues are colored according to their codons.

Multiple residues of the nascent peptide, which is largely contained within the ribosome at takeoff, interact with the ribosome’s peptide exit tunnel, causing a slowing of translation and a hyper-rotation of the two ribosomal subunits. These interactions contribute to dissociation of the peptidyl-tRNA from its codon and retention of the peptidyl-tRNA within the ribosome after takeoff.

A key requirement for bypassing is the potential to form a stem loop (take-off SL) structure in which the take-off codon is in the 5’ side of the stem. During take-off, the top part of the SL, known as the A-site SL, transiently forms in the ribosomal A-site (aminoacyl-tRNA Acceptor site) [29, 30], and performs key roles in several aspects of the bypassing mechanism.

The take-off codon, Gly46, is followed by a UAG stop codon whose slow decoding is significant for the efficacy of take-off [17, 23]. The equivalent codon in yeast mitochondrial bypassing is a sense codon whose tRNA is not present, causing a similar slowing. Avoidance of ‘normal speed’ decoding in the ribosomal A-site, facilitates peptidyl-tRNA dissociation from the P-site.

Specific sequence 5’ of nucleotides that participate in the take-off SL, can pair to form a separate SL, the 5’ SL, that forms after its sequence exits the ribosome [28, 29, 32]. While it may have a role in blocking backwards bypassing, its main proposed function is that its formation helps propel the initial pseudotranslocation steps of bypassing [28, 29, 31, 32].

Once the peptidyl-tRNA anticodon has dissociated from the take-off codon, the tRNA retained in the P site scans just the 3’ part of the bypass mRNA for a matching Gly codon [29]. When a complementary triplet enters the P-site, the tRNA re-pairs with the mRNA at this ‘landing’ (or ‘anticodon re-pair’) site, enabling the resumption of translation at the 3’ adjacent codon. Landing is enhanced by binding of the anti-Shine Dalgarno sequence of the bypassing ribosome to an (A)GAG(A) sequence 6 nt upstream of the landing site (by analogy with the arrestor gear on an aircraft carrier). A further mRNA stem loop, the Forward Slippage Barrier (FSB), is at the leading edge of the ribosome when the peptidyl-tRNA anticodon reaches the landing site. The FSB may impede the bypassing ribosome and promote correct landing [28]. The specificity of ‘correct landing’ is high since the only in vivo bypass protein product from the T4 sequence identified by mass spectrometry is derived from landing at the GGA 50 nt 3’ of the take-off site [27].

The insert that splits *gene 39* from *gene 60* contains a pseudogene of a homing endonuclease, MobA. Related phage encode an intact MobA protein, which was shown to be active and capable of copying the region around it to a related T2 genome, horizontally transferring both the split gene and the 50 nt insert, during a mixed infection, [19]. The bypass sequence within *gene 60* then protects the target genome from further nicking by MobA. Homing endonucleases are part of mobile genetic elements, cutting DNA at specific sites in genomes that lack the element, and inserting themselves in the target genome by recombination-mediated repair [33]. T4 contains 15 such homing endonucleases [34], and related phage share many and contain others, indicating that these elements continue to spread and be lost [35]. Some are free-standing endonucleases that insert between genes while others have the endonuclease within a group I intron or flanked by intein elements, which allows them to insert within ORFs without disrupting them. The mobA insertion has neither safeguard. It both splits the primary ORF as well as introducing the bypass cassette and so requires multiple mechanisms to resolve these problems.

Despite the scale of the efforts to understand the origin and especially the mechanism of bypassing, there have been no reports of similar bypassing in other phages. We have sought, and found, novel occurrences of gene 60 bypassing. They provide insight into both the evolutionary origin and mechanistic latitude with their future experimental analysis likely to yield deep understanding of the parameters necessary for such dramatic departure from successive non-overlapping triplet decoding.

## Results

### Homologs of *gene 60* bypass are found in many phage

We searched the NCBI NR sequence database for additional cases of the Topoisomerase 2 large subunit gene (Topo2L, *gene 39*) with a disrupted ORF and the possibility of similar translational bypass to that of T4. We found 28 genomic sequences encoding a Topo2L ORF split by insertion of a MobA and predicted translational bypass cassettes (Table 1, clades 1-6). These include 14 close relatives of T4 and 13 sequences from 5 other clades of Myoviridae phage, all with gamma proteobacteria hosts: *Salmonella, Shewanella, Proteus, Acinetobacter*, and *Aeromonas*. This genomic diversity allows us to identify functional bypass cassettes and to explore their evolutionary dynamics.

**Table 1.**
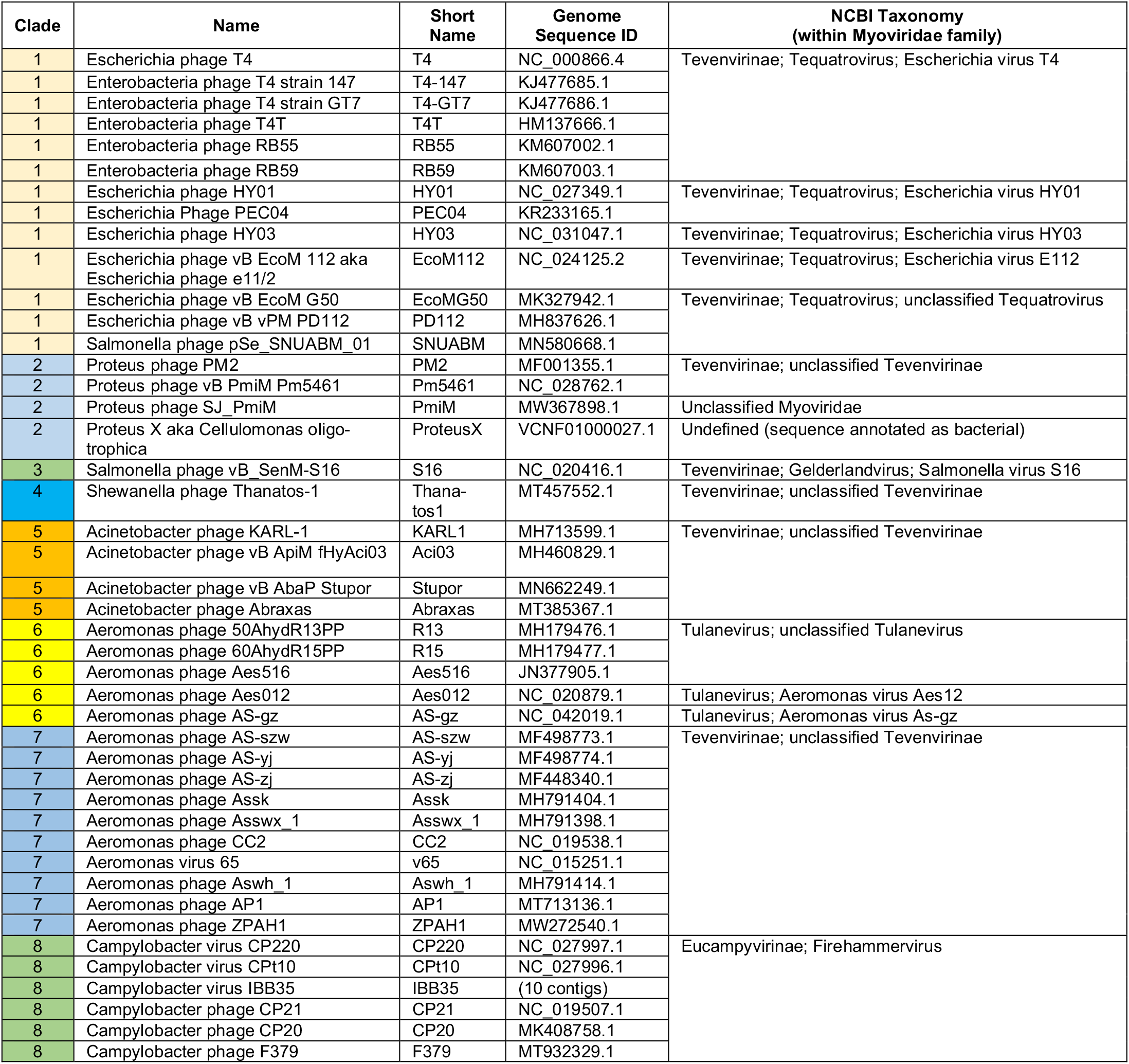
Genomes with bypass cassettes.

### The MobA element has multiple independent insertions in the same location

For all 6 clades of phage, we found genomic sequences that were highly similar (96-99% DNA sequence identity within the Topo2L gene, File S1), but lacked the split ORF and MobA element, indicating that the insertion may have happened independently in each clade. Alignment of the shared N terminus of Topo2L also shows tight pairing of species with and without the MobA insertion (Fig S1A). Alignment of MobA protein sequences shows near-identity within clades, and substantial diversity between clades, though largely reflecting the phylogeny of their hosts, suggesting that each element is recently active within a clade, and more anciently related between clades (Fig S1B).

Alignment of related genomes with and without the MobA insertion allows mapping of the insert site in all 6 clades (File S1). Each alignment shows a clean break in sequence identity on the 5’ end of the insert and at most a few nt of ambiguity on the 3’ end, and so defines the likely extent of the MobA mobile element. The element has a similar structure and insertion site in all clades (Fig 2, Table S1, File S1). It is 1097-1106 nt long, consisting mostly of an HNH-family homing endonuclease, MobA, of 269-279 AA (807-837 nt). The insertion splits the Topo2L ORF in two, always adding an in-frame novel C-terminal extension (CTE) of ∼44 AA to the upstream ORF1 (*gene 39*), and an N-terminal extension (NTE) of 44-45 AA, followed by the translational bypass cassette, to the downstream ORF2 (*gene 60*) (Fig 2, 3). The insertion causes the deletion of 55-75 nt of the Topo2L ORF, which is largely replaced by a homologous region (HomR) encoded within the NTE of the MobA insert. The homR contains the site where MobA was seen to nick and insert in phage T2 [19].

**Figure 2.**
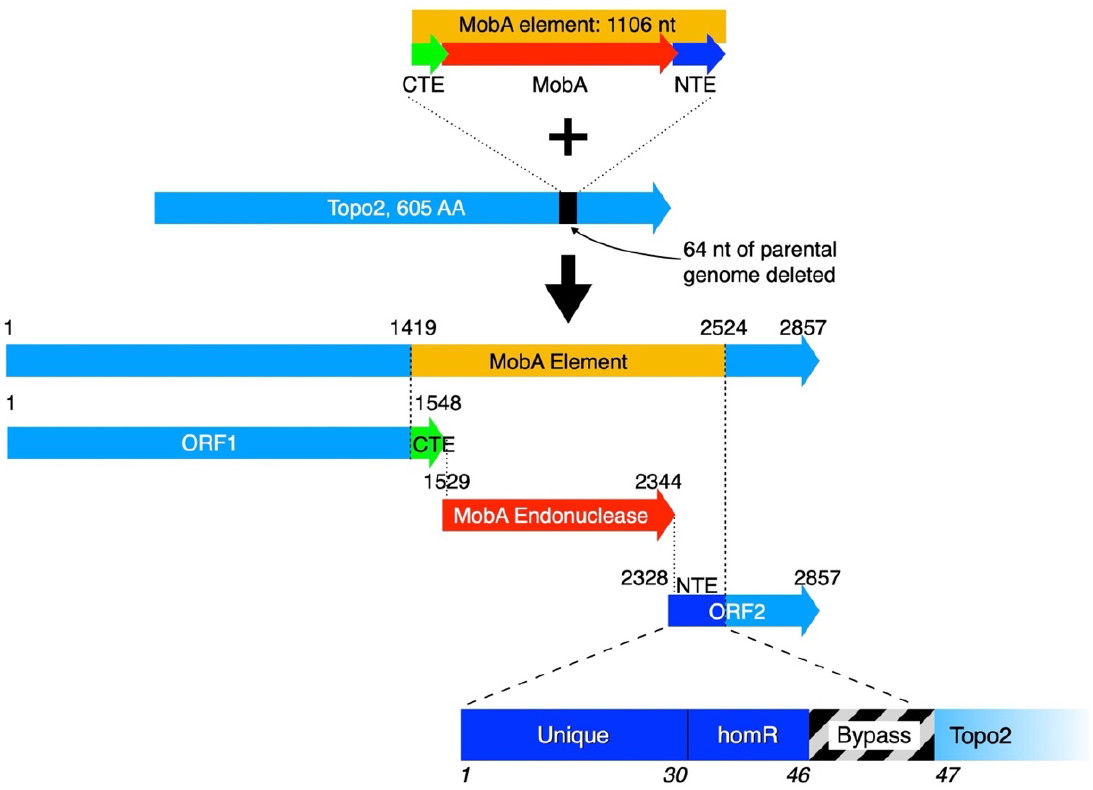
Schematic of MobA element insertion, using phage HY01 as example. The MobA mobile element, encoding the MobA endonuclease (red) flanked out-of-frame by C-terminal and N-terminal elements (CTE and NTE, green and dark blue, respectively) inserts into the Topo2L gene, deleting a 64 nt region (black). The inserted genome encodes a split Topo2L (light blue), with the 515 AA ORF1 (gene 39) containing the N-terminal portion (472 AA) plus the 43 AA CTE sequence, while ORF2 (gene 60) consists of the remainder of the parental Topo2L (117 AA), preceded by the element-encoded NTE (46 AA) and the bypass sequence (hatched). Blow-up of NTE shows that it consists of a unique 30 AA N-terminus, an insert-encoded region of 16 AA with homology to the deleted region (homR), and the 50-nt translational bypass cassette. In clade 1 but not in other clades, the MobA element contains an additional 9 nt of homR beyond the bypass cassette. Nucleotide co-ordinates labelled, relative to start of Topo2L, AA residues of ORF2 labelled in italics.

**Figure 3.**
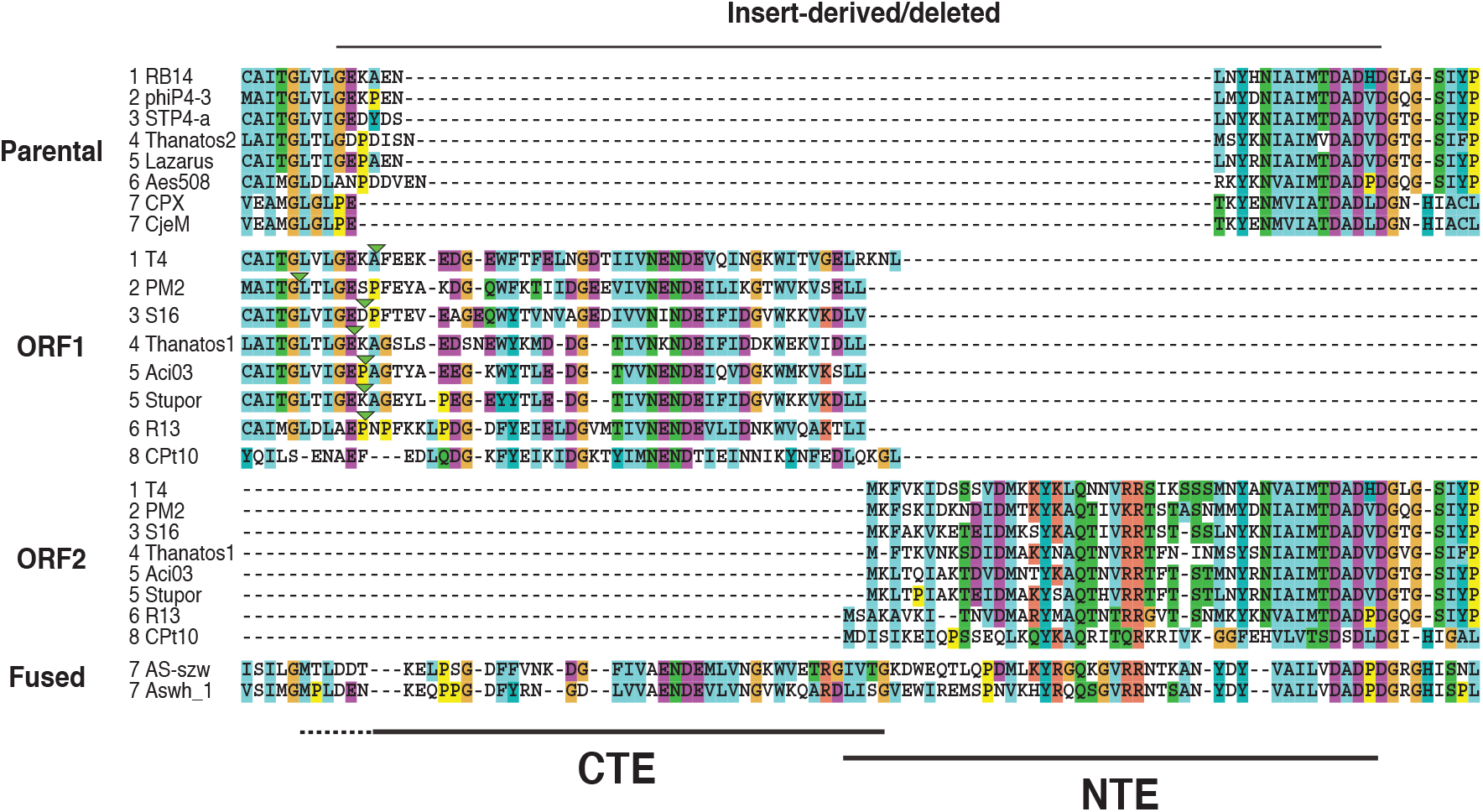
Protein alignment of topoisomerase genes. The top 8 sequences are from parental genomes (no MobA insert), followed by the split proteins, ORF1 and ORF2, and the clade 8 fused ORFs. Names are labelled with clade number. CTE and NTE are the insert-derived termini of ORF1 and ORF2. Inverted green triangles mark the 5’ end of insert sites, where known. Residues are colored by conservation and physicochemical properties using the Clustal coloring scheme.

In some genomes, including the lab T4 strain, KARL1, and Aes012, the endonuclease is pseudogenized or partially deleted, and others have entirely lost the endonuclease, while retaining the bypass cassette (Fig 4).

**Figure 4.**
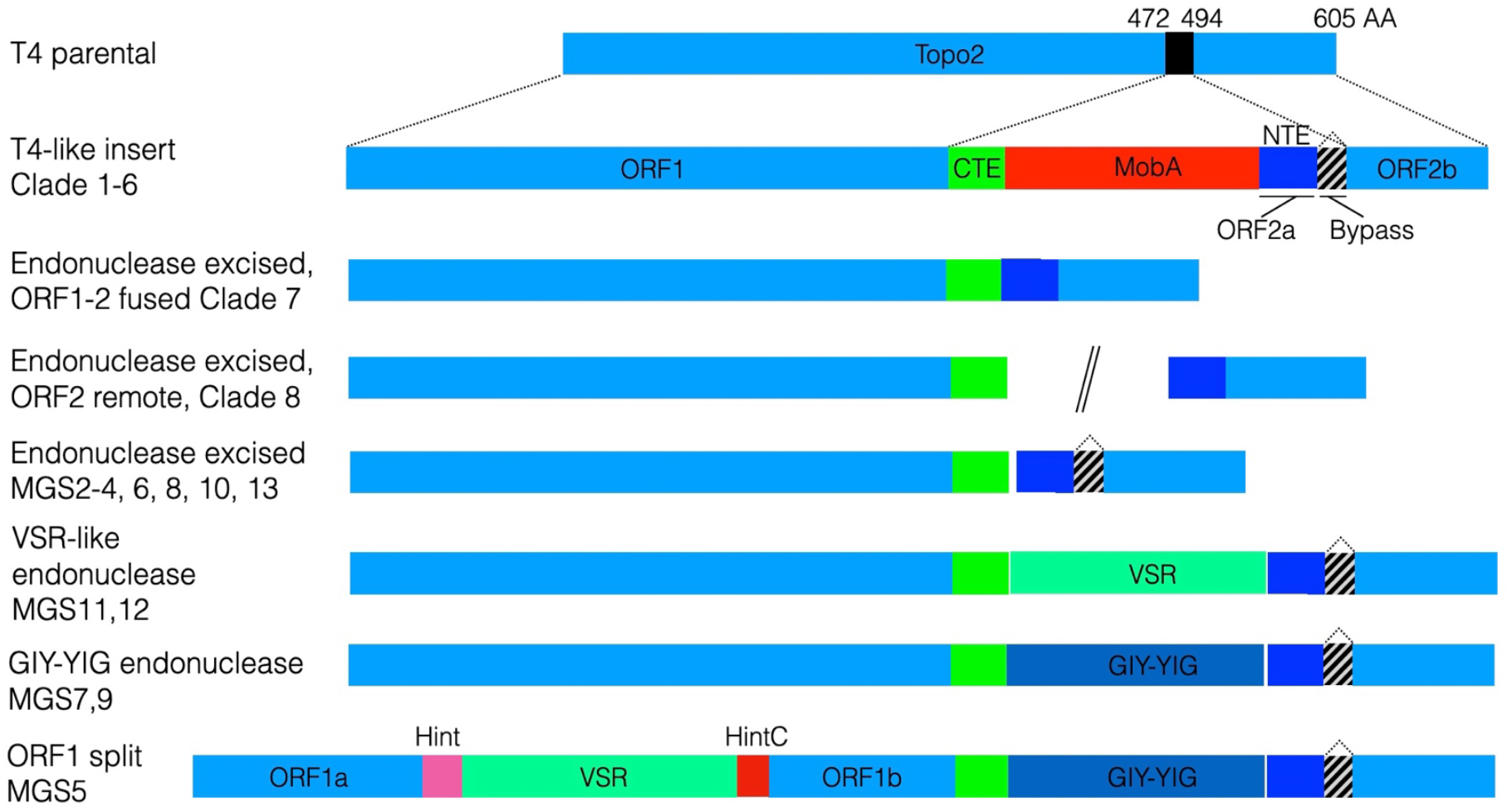
Varied gene structures of bypass insertion. MobA, GIY-YIG and VSR are structurally distinct families of homing endonucleases, Hint and HintC are intein domains. Hashed boxes denote bypass region, Colors as in Fig. 3

The insertion site varies slightly, from nt 1399-1419 of Topo2L (*E. coli* phage RB14 numbering; AA 466-473), and this variation is mirrored by the extent of CTE sequence at the start of the insertion, such that the CTE is always in-frame, and varies in length, proportional to the position of insertion (Fig 2). The CTE protein sequence is unique but selectively conserved, indicating that it has conserved function. The CTE is in a different frame to MobA, and their ORFs overlap, by 1-20 nt (Fig 3, Table S1).

The 3’ end of the MobA insertion always maps just downstream of D490 or G491 of Topo2L (*E. coli* RB14 numbering), with the exception of clade 1 and 4, where it extends 9 and 8 nt further, respectively. The NTE of ORF2 is more complicated. It starts with a ∼30 AA, partially conserved, unique sequence, followed by the ∼15 AA HomR region that is homologous to the Topo2L sequence that was deleted during insertion, and then the translational bypass cassette, encoding stops and/or frameshifts (Fig 3, Fig S2). The insert ends at the landing codon or shortly after. The NTE also overlaps, out-of-frame, with the MobA ORF, by 8-32 nt.

The insert-encoded HomR sequence is distinct from the parental sequence (67-81% DNA identity) and slightly shorter than the deleted sequence, resulting in loss of 2-7 AA relative to the parental sequence, in a region that is not highly conserved in Topo2L (Fig 3). This shows that the HomR is not derived from the parental Topo2L during insertion. However, in each clade, it is more similar in sequence to the parental Topo2L than to the homR of other clades, indicating that there is some re-copying of the homR from parental sequence within each clade, rather than being created once, early in MobA element evolution.

Two other Myoviridae clades (7 and 8, infecting *Aeromonas* and *Campylobacter*) showed evidence of MobA-mediated splitting of the Topo2L ORF. In both cases, the CTE and NTE sequences survive, but both the endonuclease and the bypass cassette have been lost (Fig. 4).

### The *Aeromonas* clade shows evidence of hit and run

A set of 8 *Aeromonas* phage (Table 1, clade 7) show evidence of MobA insertion followed by loss and then recreation of a single Topo2L ORF. All 8 have a full length Topo2L ORF with a unique insertion that encodes CTE- and NTE-homologous sequences at the usual MobA insertion site (Fig 3, Fig S3). The endonuclease and the bypass cassette have apparently been precisely excised. The retention of the NTE and CTE sequences suggests that this is an evolutionarily recent event or that these sequences have some function in the fused protein.

### *Campylobacter*: Split gene with no bypass cassette

A set of 5 phage, from the Firehammervirus genus, infect *Campylobacter* bacteria, and all have a truncated ORF1 with a CTE that is similar to that encoded by the MobA insertion in T4 (Clade 8, Table 1, Fig 3-4, Fig S2). However, they have no neighboring endonuclease or ORF2. Instead, a divergent ORF2-like protein is encoded elsewhere in the genome, with sequence similarity to both NTE and Topo2L. (Table 1, Fig 4, File S2). These data suggest that a MobA inserted into the Firehammervirus common ancestor, then later excised, leaving the NTE and CTE sequences, with subsequent loss of the bypass region and translocation of ORF2 to another location. The continued presence and sequence conservation of the CTE and NTE indicate that they may be important for the function of the split topoisomerase.

### Bypass cassettes from metagenomic sequences

We searched for additional cases of bypassing in 18,472 metagenomic sequencing projects from the Sequence Read Archive (SRA). We found 3 candidate cases of translational bypass, MGS1-3 (Table S2), and additional cases where a CTE or NTE sequence indicated the previous presence of a MobA insertion, but where no bypass sequence was seen. We found another 10 unique cases of bypassing (MGS4-13) in metagenomic assemblies from the IMG database [36] (Table S2). These were diverse in sequence but all were most similar to the *Campylobacter* clade. Notably, sequences in clades 9 and 10 had replaced MobA with a structurally distinct endonuclease, from the GIY-YIG family (Fig 4, File S6), or had a clean excision of the endonuclease. ORF1 from MGS5 was also split by a homing intein, consisting of another endonuclease, VSR, flanked by Hint and HintC domains that allow protein splicing of the interrupted ORF1. In clade 11, MobA was replaced by yet another endonuclease, from the VSR family. This raises the intriguing possibility that one endonuclease inserted and replaced another, which would be an example of a parasitic element invading a parasitic element within a parasitic phage of a bacterium.

### Sequence conservation illuminates mechanisms of by-passing

These diverse examples of bypassing allow us to examine the bypass cassettes for conserved, functional motifs, and map current models of mechanism to the evolution of the element. The major elements revealed by patterns of conservation are: All bypass sequences are inserted in the same position (+/-1 codon) of ORF2 and have paired takeoff and landing codons, encoding either glycine (G46 in T4), or the immediately upstream aspartate (D45) in the *Acinetobacter* and *Aeromonas* clades and most metagenomes. All clade 1 (*E. coli*) sequences conserve GGA (glycine) as both the takeoff and landing codon, while Thanatos uses GGU. All *Aeromonas* and most metagenomic sequences have GAC while MGS13 and MGS11 use GAU, both Asp codons. In the other cases, the third base differs between take-off and landing codons, but is always an allowed wobble position that is decoded by the same tRNA: GGG/GGA in Proteus, GAU/GAC in KARL and Aci03, and GGU/GGC in Stupor, Abraxas, S16, MGS1, and MGS12.

Most sequences have a stop codon immediately after the takeoff codon (red in Fig 5), usually UAG, but UGA in three metagenomic sequences. Exceptions are Stupor, Abraxas, and MGS1, from clade 5, whose UAG is out of frame, and so does not encode a stop. All three have the rare Arg codon AGG 3’ of the take-off codon, which is very rare in *Acinetobacter*, the natural host of this phage, accounting for just 0.08% of codons, compared with 0.4% in *E. coli* [37]. This ‘hungry codon’ [9] may cause a pausing of translation that mimics the effect of the stop codon. These three bypass sequences have in-frame stops and shift frame, also ensuring that the non-bypassed protein product is truncated.

**Figure 5.**
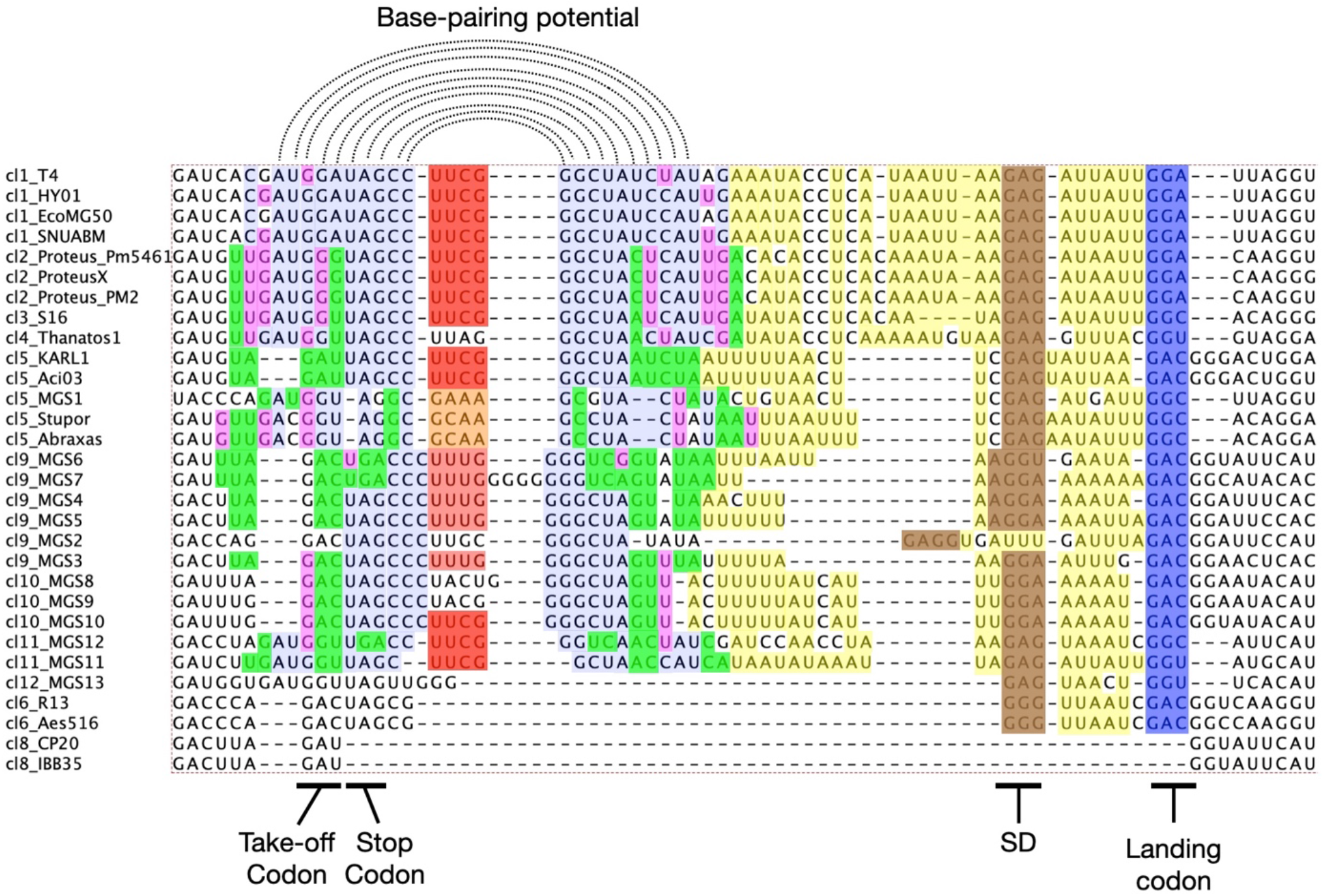
Bypass cassette alignment of non-redundant sequences. Shine-Dalgarno (SD) motif in brown, landing codon in blue. Likely base pairing of a stem-loop that includes the take-off codon is shown in light blue where identical to the clade 1 consensus, in green for W-C pairs that differ from the consensus, and in magenta for G•U or U•G wobble base pairs [65]. The UUCG and UUUG tetraloops are in dark and light red respectively, and GNRA in light orange. A and U are in light yellow in the AU-rich region between the stem loop and the landing codon.

The region surrounding the takeoff codon in T4 can form a 28-base stem-loop (SL) RNA structure, called the take-off SL. This structure is selectively conserved in other sequences, with most of the sequence changes maintaining base-pairing of the stem (Fig 5: light blue on residues identical to T4, green on substitutions that retain W-C base-pairing, magenta for allowed wobble base pairing). The tip of the SL is largely conserved as CC-UUCG-GG, a well-studied motif that is recurrent in T4 hairpins and known to stabilize their structure and act as a transcriptional terminator or to stabilize RNA [38–40]. The loop is changed to GCAA in Stupor/Abraxas, and GAAA in MG1, both matching the GNRA consensus of another highly stable loop motif [41]. Thanatos1 has UUAG, found recurrently in rRNA [39]. Metagenomic clade 9 and 10 add another C/G base pair to strengthen the stem, and have diverse loop sequences, most often UUUG, a less stable loop [42]. Clade 6 (*Aeromonas*) sequences have a drastically reduced SL sequence that may pair with downstream sequence around the SD site (GGUUA). Some of the candidate stem loops have different types of loops, meriting future investigation to ascertain the diversity of mRNA structures in the ribosomal A-site.

The T4 landing site is 6 nt downstream from a GAG sequence, which pairs with the anti-Shine Dalgarno (SD) sequence in the 16S rRNA of bypassing ribosomes (Fig 1,5). This ‘mini-SD’ facilitates peptidyl-tRNA anticodon pairing to the landing site [27]. GGG, GGA and AGG have similar pairing potential, and all four triplets are seen 5-7 nt 5’ of the landing site in most other clades. The only exception is MGS2, which lacks a mini-SD at that position but whose upstream has a stronger SD sequence, GAGG. Binding of the anti-SD to this site would be retained through several cycles of pseudotranslocation, until landing, and would preclude any other site from binding the anti-SD [7, 43].

T4 promoters typically have 6-7 nt of AU-rich sequence between the SD and the AUG codon, and additional AU-rich sequence just upstream [39]. Across the range of bypass cassettes, we see a consistently similar spacing and AU-richness, in the absence of strong sequence conservation (A/U is in light green in Fig 5).

Conservation within the NTE protein sequence may indicate residues that mediate bypassing by interacting with the ribosomal exit tunnel. Many but not all of the residues implicated by cryoEM [30] or mutagenesis [28, 29] are selectively conserved (Fig S2), providing an independent perspective on their function and suggesting some variations in mechanism in different phage. Of the ribosome-interacting residues, Y16 is invariant and T40 is conserved as T or S in all bypass sequences. R24 is R or K in all but MGS11, Y33 is Y or F in all but MGS12, and K5 and L6 are also moderately conserved. M39 is partially conserved, but H44 is found only in clade 1, and so has at best a restricted function. Q19 and V22 are implicated only by mutagenesis but are highly conserved. Conversely, hydrophobic mutations to S25, K27 and S28 greatly reduce bypassing, but are not conserved. In phage that have lost the bypass cassette but kept the NTE (clade 7, 8, and an additional clade 20), there is selective loss of several of these residues (Fig S2), further supporting their role in bypassing. Y16, Q19, V22 and M39 are lost in clade 20, T40 in clade 7, and V22 and M39 in clade 8.

The T4 5’ SL, is upstream of the take-off SL, in a region of invariant sequence in clade 1, and so the SL is also conserved (Fig S4). However, this specific structure is not seen in any other clade. Examination of the best predicted secondary structure in each sequence shows substantial divergence between and even within, clades (Fig S4), but does show a consistent pattern of being somewhat stronger (lower free energy) in MobA insert sequences than the parental genomes, suggesting that some secondary structure may be functional in most clades, though not well conserved, and may not be functional in some clades, like 5 and 11, where it is particularly weak.

A third SL, the Forward Slippage Barrier (FSB), downstream of the landing site, is significant for bypassing in T4 [28]. This is encoded by the host phage rather than the insert and may have a role in Topo2L expression, but also may slow the ribosome and minimize overshooting of the landing site. As with the 5’ SL, its sequence is poorly conserved within and between clades, which have a variety of predicted secondary structures, some as strong as T4, and some, such as clades 10 and 11 being substantially weaker (Fig S5).

Each of these conserved features also shows substantial variation, suggesting some variability in function. Most dramatically, clade 6 (*Aeromonas*) has greatly reduced the take-off SL and has a much shorter bypass region of just 20 nt, compared with 50 nt in T4. Other moderately conserved residues may also be functional and locally conserved within their individual hosts. For instance, there are only 6 nt changes between the *E. coli* and *Proteus* clades within the 50 nt bypass cassette, two of which conserve base-pairing in the SL, yet the synonymous substitution rate across a collection of clade 1 and clade 2 ORFs is ∼0.4, indicating that many or most of the residues across the bypass cassette are under evolutionary constraint.

## Discussion

These genomic data show that a previously unique programmed bypassing mechanism is both widespread and variable enough to cast light on its origin and mechanistic versatility.

### Structure, functions and evolution of the MobA element

All MobA elements share several characteristics: They are exclusively inserted into the gene encoding Topo2L, splitting the encoded protein at the same location and adding conserved NTE and CTE sequences, along with the translational bypass cassette. Most MobA insertions are relatively recent events. In a mixed infection model, MobA is highly efficient at transforming phage lacking MobA, showing that it is still an active homing endonuclease [19]. The endonuclease and bypass cassette may once have been independent sequences, but it is clear in all species that they now form a single mobile element.

The conserved insert location and sequences of CTE and NTE suggest a specific mechanism to retain Topo2L function. MobA always inserts within the Toprim domain of Topo2L, splitting a key metal binding site between the two ORFs, likely making each ORF non-functional. However, ORF1 and ORF2 encoded proteins can complex with gp52 to make a functional Topo2 holoenzyme [44]. All split ORFs retain CTE and NTE sequences, indicating they may be required for dimerization or function. A similar case is seen with the related MobE endonuclease. It typically inserts between the ribonucleotide reductase genes *nrdA* and *nrdB*, but in phage Aeh1 and several others, a variant MobE inserts within the *nrdA* ORF. As seen in MobA, insertion adds CTE and NTE sequences to the split ORF products and the two proteins form a stable protein complex with each other and with NrdB [45]. Deletion of the CTE and NTE did not block binding of the split proteins to each other, but did impair their binding to NrdB and abolished enzyme activity [46]. Several other Myoviridae have independent but similar MobE insertions with conserved NTE and CTE sequences (GM, unpublished). Other homing endonucleases that insert within ORFs are flanked by intein sequences or group I self-splicing introns. The creation of CTE and NTE tails to restore function to split ORFs may be another general strategy to avoid the damaging effects of splitting an ORF. The NTE sequence also has a critical role in the translational bypass mechanism (below).

The translational bypass cassette remains unique and its biological functions a puzzle. It is found in every phage that has the endonuclease, and several that have lost it (Fig 4). Insertion disrupts the MobA recognition site on DNA and so prevents self-cleavage of the insert by MobA [19]. The nick site is near the middle of the insert-derived homR region, with 28-36 bases on either side that are divergent from the equivalent region of Topo2L. At first sight, these polymorphisms would seem to disrupt the cleavage site, yet these polymorphisms alone could not block MobA nicking, but the bypass insert, 20 bases downstream, was successful [19]. Hence, the bypass, while distal to the nick site, protects an infected phage from further nicking by insert-encoded MobA. This region of Topo2L is highly conserved, so the bypass cassette allows for the insertion of exogenous DNA without causing a potentially deleterious insertion within the protein sequence.

Such a complex mechanism to avoid MobA nicking, and the diverse distribution of MobA elements suggest that this may not be the only function of bypassing. Bypassing efficiency is in the order of 50%, which would appear costly to the phage [20, 21, 23]. Deletion of sequence between the take-off SL and the mini-SD, can increase efficiency [28], raising the possibility limited or regulated bypassing may be optimized for MobA survival, or that ORF1 alone may have some secondary function. It is also possible that the MobA element is mildly deleterious (parasitic) to its host, with its rate of deletion being matched by its rate of spread in mixed infections. This might explain how most insertions appear to be evolutionarily recent, even though the element itself is far older.

We see progressive loss of the MobA element. The endonuclease has been pseudogenized in T4, Karl-1, and Aes012, and several other sequences have lost the entire endonuclease, while retaining the bypass (Fig 4). Others have also lost the bypass, while retaining the split Topo2L and CTE/NTE tails (Clade 7; Clade 20 in Table S2) or undergone secondary fusion (Clade 8). Thus, the endonuclease is no longer essential after insertion, and the bypass may not be needed after endonuclease loss, but the split Topo2L always retain the CTE and NTE sequences.

The replacement of MobA by VSR or GIY-YIG endonucleases in some metagenomic sequences shows that the bypass cassette is also independent of the mobilizing endonuclease. These shifts have some precedent. T4 has a related MobC inserted between *nrdD* and *nrdG* genes, and in some related genomes, MobC is replaced by a GIY-YIG gene, *segH*, while MobE (between *nrdA* and *nrdB*) has been replaced by a VSR-like endonuclease [47]. It is curious that these endonucleases appear to have evolved a very similar selective DNA target site, which may reflect the very high sequence divergence in these gene families.

### Mechanism of bypassing

The diversity of bypass sequences shows clear selective conservation of elements known to be functional for T4 bypassing as well as novel variations whose future detailed study may reveal additional mechanistic latitude. Sequence changes in the takeoff and landing codons explore the multiple variations that allow a single tRNA to bind to both codons; most of the variations in the take-off SL maintain basepairing in the stem, and while the mini-SD has substantial variation in sequence, it selectively maintains the core and AU-rich flanking regions. Some of these elements break down in more divergent clades, including substitution of non-homologous stem-loops, and the swapping of a stop codon for a rare ‘hungry codon’, and possible loss of some entire elements, and likely represent mechanistic variants of the same overall theme.

### Nascent peptide decoding & exit tunnel interaction

Interactions between the nascent peptide and the ribosome have major effects on ribosome slowing, ribosome conformation at take-off, codon: anticodon dissociation, retention of peptidyl-tRNA during bypassing and an influence on landing site fidelity. For the conditions tested, cryoEM has revealed clear interactions with specific residues and assays with mutants have shown functional significance; the newly identified sequence diversity now adds a complementary perspective. In the unique region (AA 1-31), the most conserved residues are supported by mutagenesis or structure analysis, though the reverse is not always true: Mutagenesis and cryoEM showed a role for S28, but this position is weakly conserved. Mutation of nearby non-conserved residues S25 and K27 to hydrophobic amino acids also reduced bypassing; these are all weakly conserved as polar, so it may be that the hydrophobic substitutions are specifically deleterious.

Within the homR region (AA 32-46), the protein sequence is constrained by Topo2L function, and many residues are more similar to their parental sequences than to other inserts, with no clear signal of convergent evolution to support bypassing. For instance, H44 which was suggested to block rotation of the ribosome [30] is conserved in clade 1 insert and parental genomes, but is not seen in any other clades. Other residues may function both in Topo2L and bypassing: D45 is part of the conserved TOPRIM motif, and also shown to diminish release factor recognition at A-site stop codons [48] and has a positive effect on bypassing [23].

Some of the conserved residues not seen as contact points in the cryoEM visualization of ribosomes paused at take-off, such as Q19 and V22 may be involved in the initial slowing of the ribosome prior to takeoff, as seen by smFRET [29]. The more unusual variants may be valuable for experimental analysis to understand the sequence dependence of the slowing, and to resolve puzzling cases such as MGS13, whose 6 AA insertion appears to position the residues involved in slowing well before the take-off codon.

### Poor A-site decoding potential

When the take-off codon is in the P-site, the potential for the A-site codon to be quickly decoded is exceptionally low. This facilitates A-site mRNA structure formation and P-site anticodon: codon dissociation. The nature of the A-site codon (UAG in T4 and most other sequences) and of its flanking sequence, are among the features relevant to its low coding propensity. Three sequences have an alternative stop codon, UGA, which in T4 gives a modest increase in bypassing efficiency [23]. In some clade 5 sequences, the A-site codon is the ‘hungry’ sense codon AGG, which constitutes only 0.08% of codons in Acinetobacter, the natural host of these phages. This indicates that the cognate tRNA is similarly rare and predicts relevance of very slow decoding, consistent with results from frameshifting [25, 49], non-programmed bypassing [50] and yeast mitochondrial byp bypassing [13].

### Matched take-off and landing sites

All bypass sequences are inserted at the Gly46 or Asp45 codons, both invariant residues in Topo2L. The T4 take-off and landing codons are both GGA, which binds tRNA2Gly [22], whose anticodon is ^5’^mnm^5^UCC ^3’^ and has the potential to also pair with GGG. The tRNA is retained in the bypassing ribosome after takeoff, and then binds again to the landing codon. Re-pairing to mRNA occurs in the ribosomal P-site where anticodon: codon pairing is not monitored as in the ribosomal A-site for incoming aminoacyl-tRNA. Though influenced by the restraint provided by mini-SD pairing and with consequences for efficiency, considerable latitude has been detected [51]. Even without this consideration, the conservation pattern in the newly identified occurrences is strongly consistent with re-pairing of the retained peptidyl-tRNA to matched landing sites. When takeoff is at codon G46, the takeoff and landing codons are either identical, or have third codon base wobble pairing potential, either G/A or U/C, such that the same tRNA^Gly^ can pair with both the take-off and landing sites. When takeoff has shifted to Asp45, GAU and GAC are always used for takeoff and landing codons, both of which can be paired with by the same tRNA^Asp^. A similar tolerance of wobble bases and even larger deviations was seen in yeast mitochondrial bypassing [13]. The 3’ parts of the bypass region are always selectively devoid of triplets that could act as premature landing sites.

### mRNA:rRNA pairing at the coding gap SD

T4 uses a mini-SD, GAG, to facilitate pairing of the retained peptidyl-tRNA with the landing site codon 6 nt 3’ [27]. With the exception of MGS2, all sequences have a mini-SD within 1 nt of that site. Variations in the G-rich mini-SD reflect a need to avoid premature landing sites, as many phage with a Gly46 take-off codon have a GAG mini-SD, which is a possible landing codon for those with Asp45 take-off, while the mini-SD of some sequences with Asp45 takeoff (e.g. GGA), is a possible landing codon for Gly46 take-offs.

When the landing site has strong anticodon pairing potential, the stimulatory effect of the mini-SD is modest, but for cases with weaker pairing potential, such as the GAU in Karl1 and Aci03, the effect is likely greater.

MGS6 (clade 9) differs slightly from the others as its mini-SD is AGG (cf. [52]) rather than GAG or GGA and is 1 nt 5’ of nearly all others. The bigger exception is MGS2 (Clade 9) where instead of a 3 nt mini-SD, a stronger SD, GAGG, is seen, and it is 12 nt 5’ of the landing site. Based on studies with translating ribosomes [43], at this spacing the rRNA:mRNA pairing should be intact and remain so for an additional cycle or two, but the increased tension has a tendency to ‘pull back’ (i.e. 5’ with respect to the mRNA) anticodon pairing by 1 nt. Extrapolating from translating to bypassing ribosomes, rRNA pairing to the SD in MGS2 would preclude pairing with a mini-SD nearby 3’ because of retention of the original pairing [7], and so it is not surprising that a mini-SD at the ‘standard’ position is absent in MGS2. In R13 (clade 6) the mini-SD is only 5 nt 3’ of the take-off site, and 3 of these are the stop codon 3’ adjacent to the take-off site. Though rRNA anti SD scanning and pairing are expected, peptidyl-tRNA anticodon scanning in this case before the landing site is likely minimal at most.

### AU-richness in the 3’ part of the gap

Beyond the take-off SL, the rest of the bypass region is markedly AU-rich, though the specific sequence is not conserved (Fig 5). This may enhance SD function, since it is typically flanked by AU-rich sequence when found at the start of T4 genes [39], and the spacing of 5-6 nt between mini-SD and landing codon is similar to that between SDs and translational starts. This may also reduce the incidence of G-C driven secondary structure that might impede ribosomal sliding, as detected experimentally [28]. *M. capitatus* byp bypassing does not use an SD (these are not used at all in these yeasts) but the published sequences [13] also show a striking AU-enrichment in the 3’ part of almost all byps.

### 5’ SL and Forward Slippage Barrier

In contrast to the selective conservation of the take-off SL, the 5’ SL and FSB structures are not conserved between clades, having a wide variety of predicted secondary structures in different sequences, many of which are of similar strength to randomly selected fragments of T4. Though RNA structures predictions are imperfect [53, 54], the wide diversity of predicted structures highlights that these two structures are much less conserved than other functional elements within the sequence. It may be that these T4 structures are simply fortunate accidents that increase experimental bypass rates, and are not important for survival. Since the FSB sequence is from the host phage, and is conserved in parental sequences, it clearly has not evolved for a bypassing function. Indeed, when Samatova discovered the FSB, it was suggested that it may not be sequence specific on the basis of complex results from mutation experiments [28]. There may be conservation of the potential for continuous pairing of mRNA directly on its exit from the ribosome, which may involve a set of less stable structures that form and break during translocation, rather than a discrete 5’SL. This would not be seen in our static secondary structure analysis. It is also possible that there is genuine variation in the strength of these elements, but that they are either compensated by other bypass functional elements or that the degree of bypass is distinct in different phage. While we cannot conclude that lack of conservation implies lack of function, the sequence diversity seen will allow experimental testing of biologically relevant variants.

## Materials and Methods

New bypass sequences were identified by Blast [55] searches against the NCBI NRAA database, the JGI IMG database [36] and the Sequence Read Archive (SRA) [56]. Initial searches of NRAA with the first 60 AA of T4 gene 60 and the last 60 AA of gene 39, against NRAA used a cutoff of e<1e-4. Matches were manually inspected for NTE and CTE sequences, the surrounding nucleotide sequences were extracted from GenBank and manually inspected for presence of a bypass region. Blast2 was used to detect presence of a MobA gene or pseudogene. This search was repeated with example sequences from each additional clade, and profile HMMs for the CTE and NTE were searched against NRAA and NRNT, using HMMER3 [57]. SRA searches were carried out in Jan-May of 2021 against approximately 14,000 datasets annotated as metagenomic or viral, using Amazon Cloud servers. Additional searches using full length Topo2L to detect any truncated ORFs failed to find any that could not be explained by poor sequence quality or poor annotation.

Parental genomic sequences were identified using BlastN with a query covering 200 nt on either side of the insert site and aligned manually. Divergent VSR and GIY-YIG endonuclease sequences were detected with psi-blast and by alignment to custom profile HMMs Sequences were aligned with Clustal Omega [58] or muscle [59] and hand-edited in JalView [60]. Neighbor-joining trees were created in PhyML 3.0 [61] with default parameters and visualized in FigTree 1.4.4 [62] and Adobe Illustrator. RNA secondary structure was predicted with RNAfold [63], and DNA sequences were translated with Expasy [64]. Alignment figures were rendered in JalView using the Clustal color scheme, and were annotated in Adobe Illustrator.

## Authors’ Contributions

Following contact from JFA, both authors initiated the project. All the bioinformatics analysis was performed by GM, with JFA integrating the results with the mechanistic aspects. Both authors wrote the manuscript.

## Acknowledgements

We thank J.D. Puglisi for an earlier SD insight, Wayne Matten of NCBI for help in setting up SRA searches, and D.J. McConnell for also prompting a repeat of earlier bioinformatics searches and support. This work is dedicated to the memory of great interactions with the late Alan J. Herr with whom JFA will have no more stair races or banter.

## Supplemental Online Material

**File S1**. FastA alignments of genomic regions of phage with and without the MobA insert, clades 1-5 (.zip).

**File S2**. FastA alignment of Topo2L sequences, including ORF1, ORF2, parental, and read-through sequences from complete genomes and metagenomic sources.

**File S3**. Bypass region, FastA alignment.

**File S4**. HNH Endonuclease protein sequence alignment. MobE, MobZ and MobY1 are the closest homologs of MobA, but have little or no sequence similarity in the C-terminal region. MobE_split sequences are inserted within the *nrdA* ORF.

**File S5**. VSR-like endonucleases aligned with homologs from NRAA and from JGI IMG database

**File S6**. GIY-YIG-like endonucleases aligned with homologs

**Supplemental Text**. Sequence analysis of endonuclease proteins.

**Table S1.**
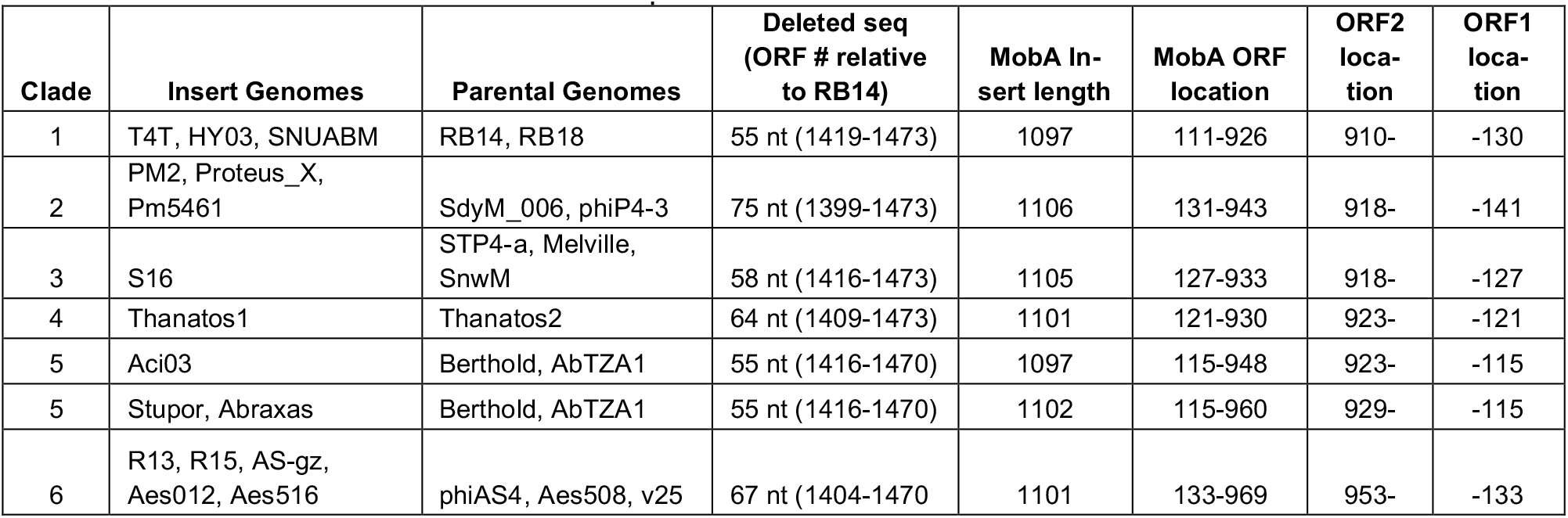
Insertion co-ordinates of MobA element across different clades. Genomic sequence alignments of phage with and without MobA insertions (File S1) were used to identify insertion points and to map ORF co-ordinates. The lab T4 strain has inserts relative to WT and is not represented here. Nucleotide co-ordinates are relative to the start of the insert.

**Table S2**. Metagenomic sequence annotations (Excel, separate file).

**Figure S1.**
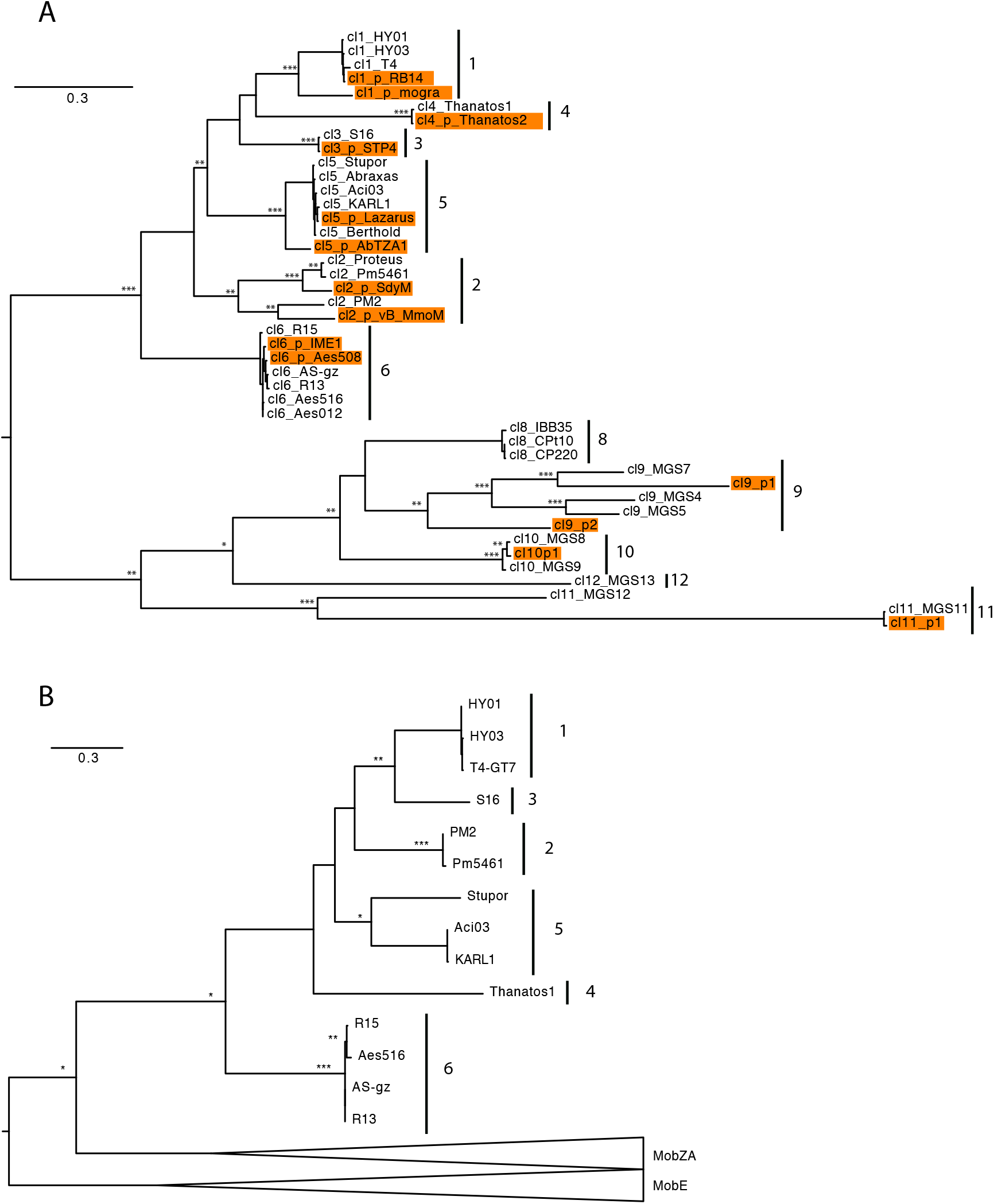
(a) Phylogenetic tree of N-terminal topoisomerase sequences, with clades numbered. Orange boxes and _p suffix show sequences from parental genomes. Nodes with high bootstrap values (from 1000 replicates) are labelled: * for >70% support, ** for >90%, *** for >99%. Pairing of insert and parental genomes indicates that independent insertions into Topo2L occurred at least 10 times. (b) Protein sequence similarity of MobA endonucleases. All MobA cluster together, distinct from their closest relatives, MobZA and MobE (see Supplemental Text). The tree reflects the phylogeny of the phage and their hosts, supporting either ancient insertion and co-evolution with the host bacterium, or more recent infections within a host clade.

**Figure S2.**
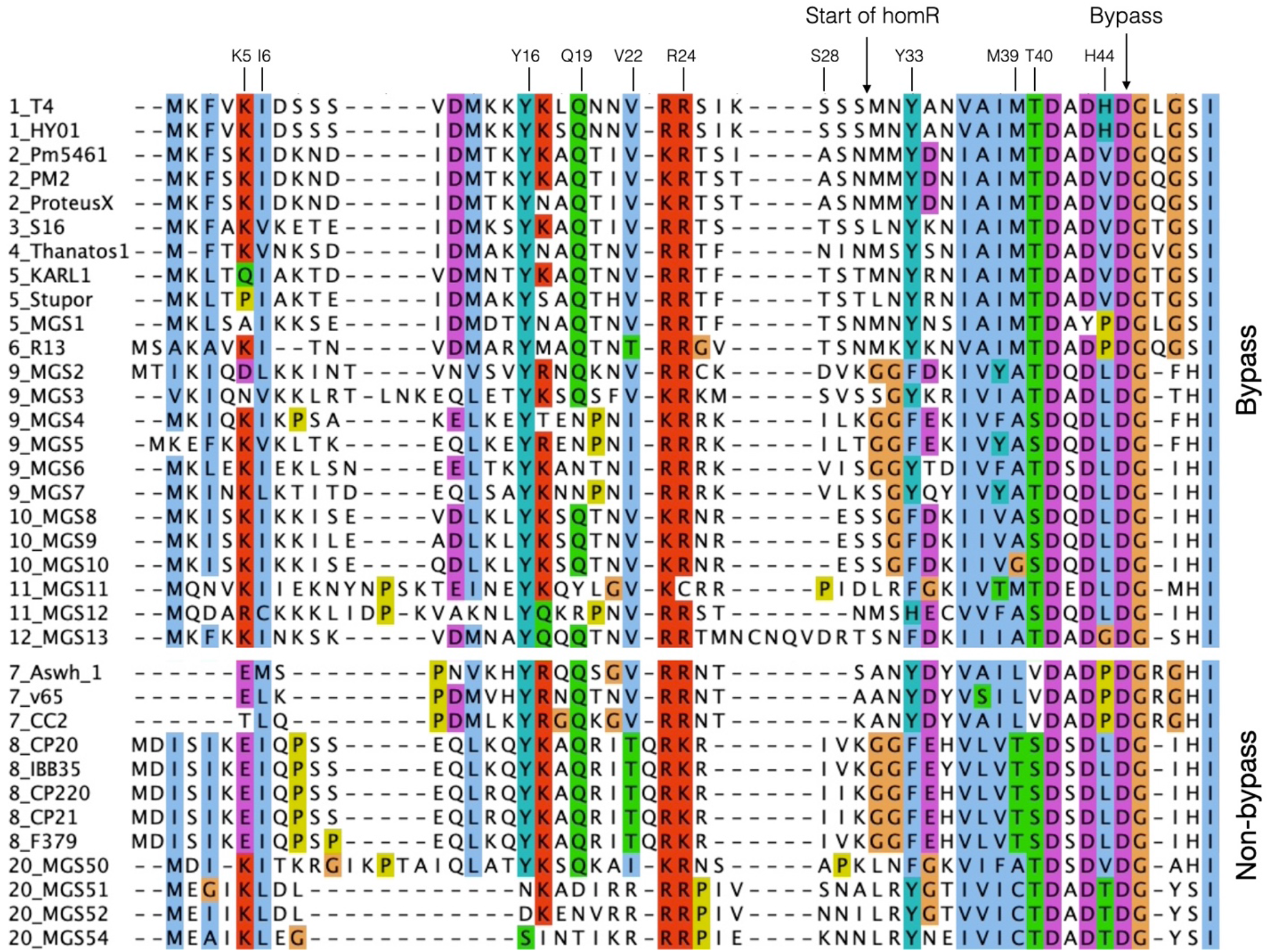
Alignment of the start of ORF2, showing non-redundant sequences. Arrows mark the start of homology with parental Topo2L genes (homR) and of the bypass region insertion. T4 residues implicated in bypassing are labelled. Lower half shows sequences from phage that have lost the bypass region and no longer need to slow translation. Residues are colored by Clustal scheme.

**Figure S3.**
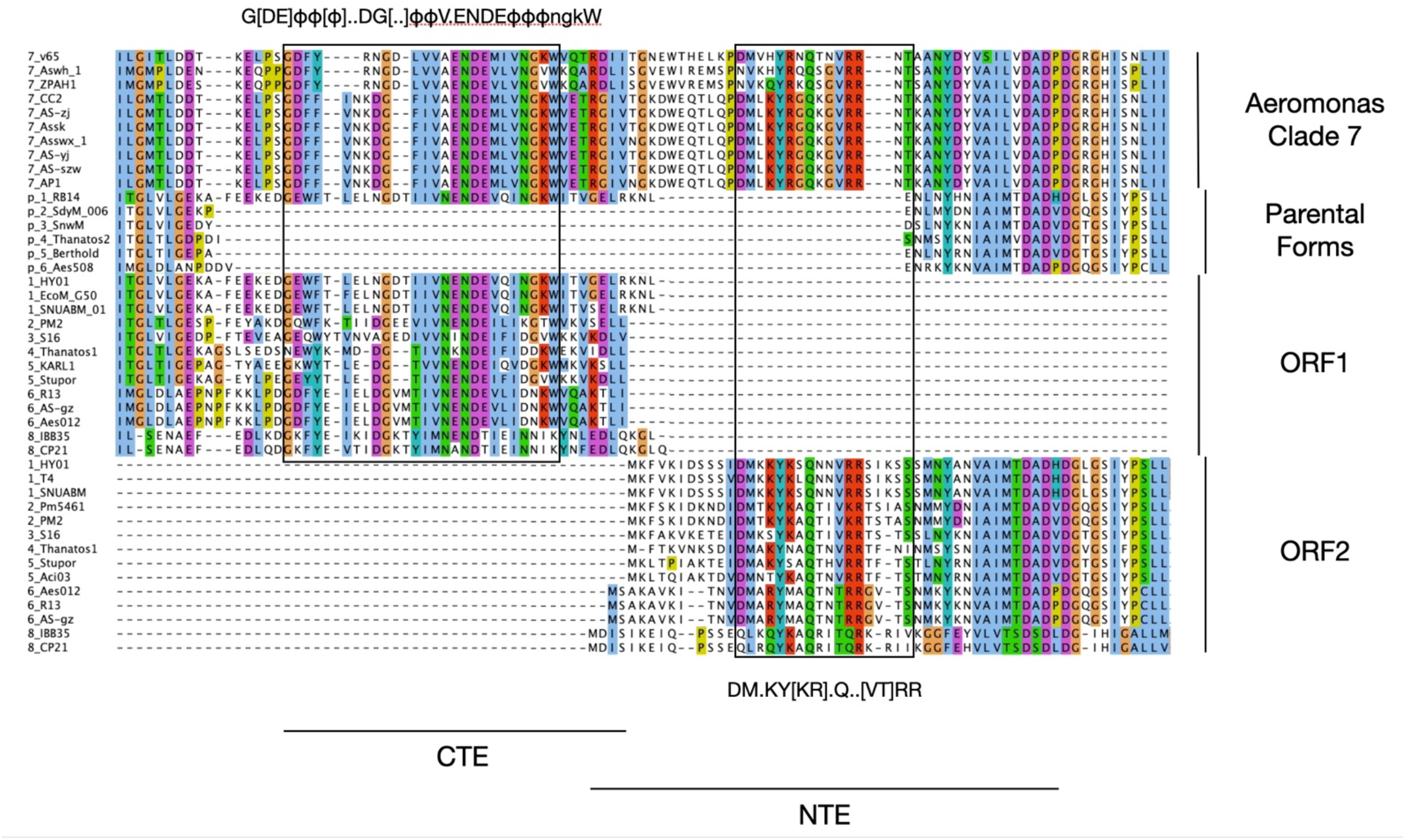
Alignment showing conservation of CTE and NTE sequences in *Aeromonas* Clade 7, where ORF1 and ORF2 are fused. Conserved CTE and NTE cores are boxed and labelled with consensus sequences. Sequences are labelled with clade number and p for parental sequences and colored by Clustal scheme.

**Figure S4.**
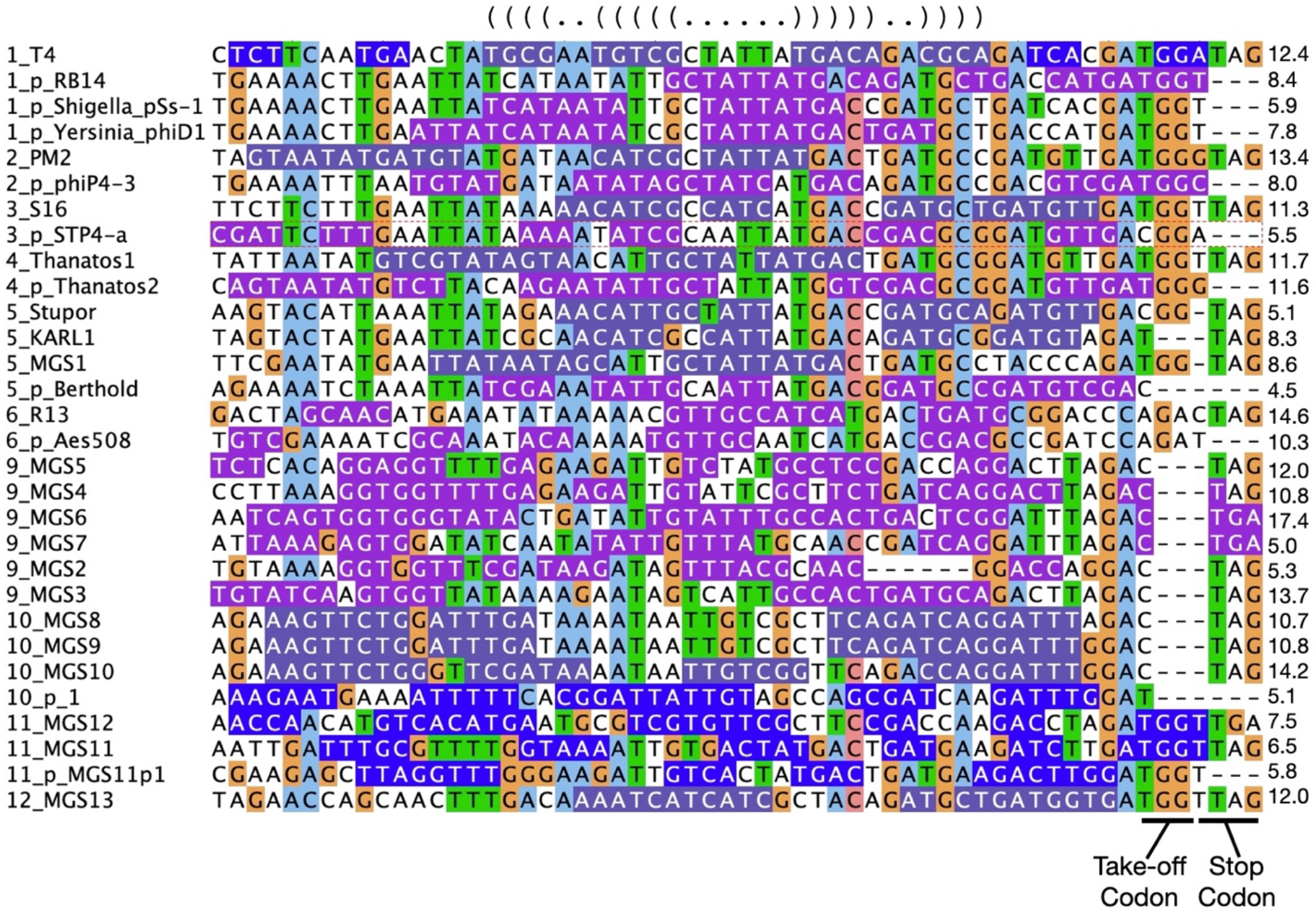
Amino acid-guided DNA alignment upstream of the take-off codon. Residues are colored by Clustal scheme, and those predicted to base pair in the optimal RNA structure predicted by RNAfold are highlighted in purple. Dot-bracket notation on top shows the structure of the T4 predicted 5’ SL. Free energy of the optimal structure (in kcal/mol) is shown on the right. Sequence names are prepended by clade number and a “p” indicates a parental sequence. Identical sequences are removed. All clades have some predicted secondary structure, but the strength, topology and location varies substantially between and even within clades, indicating that there is no shared, conserved, 5’ SL structure, even within clade 1.

**Figure S5.**
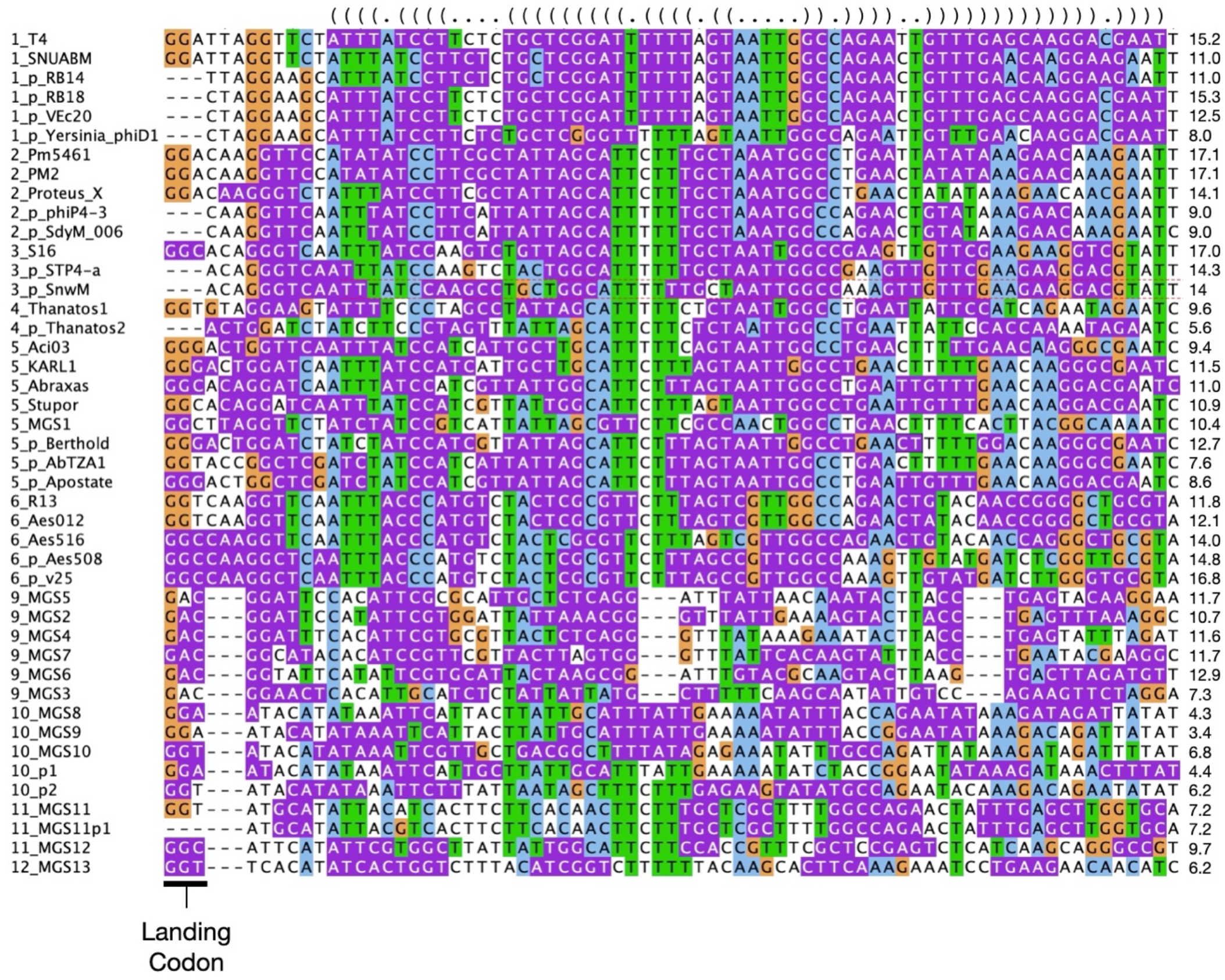
Amino acid-guided DNA alignment of the Forward Slippage Barrier (FSB) region, with base pairing of the T4 predicted FSB shown in dot-bracket notation on top. Residues are colored by Clustal scheme, and those predicted to base pair in the optimal RNA structure predicted by RNAfold are highlighted in purple. Free energy (-kcal/mol) for each optimal secondary structure is labelled on the right. Sequence names are prepended with clade number and a “p” indicates a parental sequence.

## References

1. Atkins JF, Loughran G, Bhatt PR, Firth AE, Baranov PV. Ribosomal frameshifting and transcriptional slippage: From genetic steganography and cryptography to adventitious use. Nucleic Acids Res. 2016;44:7007–78.

2. Choi J, Grosely R, Prabhakar A, Lapointe CP, Wang J, Puglisi JD. How Messenger RNA and Nascent Chain Sequences Regulate Translation Elongation. Annu Rev Biochem. 2018;87:421–49.

3. Rodnina MV, Korniy N, Klimova M, Karki P, Peng B-Z, Senyushkina T, et al. Translational recoding: canonical translation mechanisms reinterpreted. Nucleic Acids Res. 2020;48:1056–67.

4. Firth AE, Brierley I. Non-canonical translation in RNA viruses. J Gen Virol. 2012;93 Pt 7:1385–409.

5. Weiss RB, Dunn DM, Atkins JF, Gesteland RF. Slippery runs, shifty stops, backward steps, and forward hops: −2, −1, +1, +2, +5, and +6 ribosomal frameshifting. Cold Spring Harb Symp Quant Biol. 1987;52:687–93.

6. O’Connor M, Gesteland RF, Atkins JF. tRNA hopping: enhancement by an expanded anticodon. EMBO J. 1989;8:4315–23.

7. Weiss RB, Dunn DM, Atkins JF, Gesteland RF. Ribosomal frameshifting from −2 to +50 nucleotides. Prog Nucleic Acid Res Mol Biol. 1990;39:159–83.

8. Kane JF, Violand BN, Curran DF, Staten NR, Duffin KL, Bogosian G. Novel in-frame two codon translational hop during synthesis of bovine placental lactogen in a recombinant strain of Escherichia coli. Nucleic Acids Res. 1992;20:6707–12.

9. Gallant JA, Lindsley D. Ribosomes can slide over and beyond “hungry” codons, resuming protein chain elongation many nucleotides downstream. Proc Natl Acad Sci. 1998;95:13771–6.

10. Gallant J, Bonthuis P, Lindsley D, Cabellon J, Gill G, Heaton K, et al. On the role of the starved codon and the takeoff site in ribosome bypassing in Escherichia coli. J Mol Biol. 2004;342:713–24.

11. Bartok O, Pataskar A, Nagel R, Laos M, Goldfarb E, Hayoun D, et al. Antitumour immunity induces aberrant peptide presentation in melanoma. Nature. 2021;590:332–7.

12. Smith MCM, Hendrix RW, Dedrick R, Mitchell K, Ko C-C, Russell D, et al. Evolutionary relationships among actinophages and a putative adaptation for growth in Streptomyces spp. J Bacteriol. 2013;195:4924–35.

13. Lang BF, Jakubkova M, Hegedusova E, Daoud R, Forget L, Brejova B, et al. Massive programmed translational jumping in mitochondria. Proc Natl Acad Sci U S A. 2014;111:5926–31.

14. Nosek J, Tomaska L, Burger G, Lang BF. Programmed translational bypassing elements in mitochondria: structure, mobility, and evolutionary origin. Trends Genet TIG. 2015;31:187–94.

15. Brejová B, Lichancová H, Brázdovič F, Hegedűsová E, Forgáčová Jakúbková M, Hodorová V, et al. Genome sequence of the opportunistic human pathogen Magnusiomyces capitatus. Curr Genet. 2019;65:539–60.

16. Huang WM, Ao SZ, Casjens S, Orlandi R, Zeikus R, Weiss R, et al. A persistent untranslated sequence within bacteriophage T4 DNA topoisomerase gene 60. Science. 1988;239:1005–12.

17. Weiss RB, Huang WM, Dunn DM. A nascent peptide is required for ribosomal bypass of the coding gap in bacteriophage T4 gene 60. Cell. 1990;62:117–26.

18. Crick FH, Barnett L, Brenner S, Watts-Tobin RJ. General nature of the genetic code for proteins. Nature. 1961;192:1227–32.

19. Bonocora RP, Zeng Q, Abel EV, Shub DA. A homing endonuclease and the 50-nt ribosomal bypass sequence of phage T4 constitute a mobile DNA cassette. Proc Natl Acad Sci. 2011;108:16351–6.

20. Dayhuff TJ, Gesteland RF, Atkins JF. Electrophoresis, autoradiography and electroblotting of peptides: T4 gene 60 hopping. BioTechniques. 1992;13:499–502, 505.

21. Maldonado R, Herr AJ. Efficiency of T4 gene 60 translational bypassing. J Bacteriol. 1998;180:1822–30.

22. Herr AJ, Atkins JF, Gesteland RF. Mutations which alter the elbow region of tRNA2Gly reduce T4 gene 60 translational bypassing efficiency. EMBO J. 1999;18:2886–96.

23. Herr AJ, Gesteland RF, Atkins JF. One protein from two open reading frames: mechanism of a 50 nt translational bypass. EMBO J. 2000;19:2671–80.

24. Herr AJ, Wills NM, Nelson CC, Gesteland RF, Atkins JF. Drop-off during ribosome hopping. J Mol Biol. 2001;311:445–52.

25. Herr AJ, Nelson CC, Wills NM, Gesteland RF, Atkins JF. Analysis of the roles of tRNA structure, ribosomal protein L9, and the bacteriophage T4 gene 60 bypassing signals during ribosome slippage on mRNA. J Mol Biol. 2001;309:1029–48.

26. Bucklin DJ, Wills NM, Gesteland RF, Atkins JF. P-site pairing subtleties revealed by the effects of different tRNAs on programmed translational bypassing where anticodon re-pairing to mRNA is separated from dissociation. J Mol Biol. 2005;345:39–49.

27. Wills NM, O’Connor M, Nelson CC, Rettberg CC, Huang WM, Gesteland RF, et al. Translational bypassing without peptidyl-tRNA anticodon scanning of coding gap mRNA. EMBO J. 2008;27:2533–44.

28. Samatova E, Konevega AL, Wills NM, Atkins JF, Rodnina MV. High-efficiency translational bypassing of non-coding nucleotides specified by mRNA structure and nascent peptide. Nat Commun. 2014;5:4459.

29. Chen J, Coakley A, O’Connor M, Petrov A, O’Leary SE, Atkins JF, et al. Coupling of mRNA Structure Rearrangement to Ribosome Movement during Bypassing of Non-coding Regions. Cell. 2015;163:1267–80.

30. Agirrezabala X, Samatova E, Klimova M, Zamora M, Gil-Carton D, Rodnina MV, et al. Ribosome rearrangements at the onset of translational bypassing. Sci Adv. 2017;3:e1700147.

31. Klimova M, Senyushkina T, Samatova E, Peng BZ, Pearson M, Peske F, et al. EF-G-induced ribosome sliding along the noncoding mRNA. Sci Adv. 2019;5:eaaw9049.

32. O’Loughlin S, Capece MC, Klimova M, Wills NM, Coakley A, Samatova E, et al. Polysomes Bypass a 50-Nucleotide Coding Gap Less Efficiently Than Monosomes Due to Attenuation of a 5’ mRNA Stem-Loop and Enhanced Drop-off. J Mol Biol. 2020;432:4369–87.

33. Stoddard BL. Homing endonucleases from mobile group I introns: discovery to genome engineering. Mob DNA. 2014;5:7.

34. Edgell DR, Gibb EA, Belfort M. Mobile DNA elements in T4 and related phages. Virol J. 2010;7:290.

35. Petrov VM, Ratnayaka S, Nolan JM, Miller ES, Karam JD. Genomes of the T4-related bacteriophages as windows on microbial genome evolution. Virol J. 2010;7:292.

36. Roux S, Páez-Espino D, Chen I-MA, Palaniappan K, Ratner A, Chu K, et al. IMG/VR v3: an integrated ecological and evolutionary framework for interrogating genomes of uncultivated viruses. Nucleic Acids Res. 2021;49:D764–75.

37. Codon Usage Database. http://www.kazusa.jp/codon/. Accessed 30 Jun 2022.

38. Bottaro S, Nichols PJ, Vögeli B, Parrinello M, Lindorff-Larsen K. Integrating NMR and simulations reveals motions in the UUCG tetraloop. Nucleic Acids Res. 2020;48:5839–48.

39. Miller ES, Kutter E, Mosig G, Arisaka F, Kunisawa T, Rüger W. Bacteriophage T4 genome. Microbiol Mol Biol Rev MMBR. 2003;67:86–156, table of contents.

40. Woese CR, Winker S, Gutell RR. Architecture of ribosomal RNA: constraints on the sequence of “tetra-loops.” Proc Natl Acad Sci U S A. 1990;87:8467–71.

41. Heus HA, Pardi A. Structural features that give rise to the unusual stability of RNA hairpins containing GNRA loops. Science. 1991;253:191–4.

42. Tinoco I. Nucleic Acid Structures, Energetics, and Dynamics. J Phys Chem. 1996;100:13311–22.

43. Larsen B, Wills NM, Gesteland RF, Atkins JF. rRNA-mRNA base pairing stimulates a programmed −1 ribosomal frameshift. J Bacteriol. 1994;176:6842–51.

44. Seasholtz AF, Greenberg GR. Identification of bacteriophage T4 gene 60 product and a role for this protein in DNA topoisomerase. J Biol Chem. 1983;258:1221–6.

45. Friedrich NC, Torrents E, Gibb EA, Sahlin M, Sjöberg B-M, Edgell DR. Insertion of a homing endonuclease creates a genes-in-pieces ribonucleotide reductase that retains function. Proc Natl Acad Sci U S A. 2007;104:6176–81.

46. Crona M, Moffatt C, Friedrich NC, Hofer A, Sjöberg B-M, Edgell DR. Assembly of a fragmented ribonucleotide reductase by protein interaction domains derived from a mobile genetic element. Nucleic Acids Res. 2011;39:1381–9.

47. Sandegren L, Nord D, Sjöberg B-M. Seg H and Hef: two novel homing endonucleases whose genes replace the mobC and mobE genes in several T4-related phages. Nucleic Acids Res. 2005;33:6203–13.

48. Mottagui-Tabar S, Björnsson A, Isaksson LA. The second to last amino acid in the nascent peptide as a codon context determinant. EMBO J. 1994;13:249–57.

49. Gurvich OL, Baranov PV, Gesteland RF, Atkins JF. Expression levels influence ribosomal frameshifting at the tandem rare arginine codons AGG_AGG and AGA_AGA in Escherichia coli. J Bacteriol. 2005;187:4023–32.

50. Lindsley D, Gallant J, Doneanu C, Bonthuis P, Caldwell S, Fontelera A. Spontaneous ribosome bypassing in growing cells. J Mol Biol. 2005;349:261–72.

51. Herr AJ, Wills NM, Nelson CC, Gesteland RF, Atkins JF. Factors that influence selection of coding resumption sites in translational bypassing: minimal conventional peptidyl-tRNA:mRNA pairing can suffice. J Biol Chem. 2004;279:11081–7.

52. Felletti M, Romilly C, Wagner EGH, Jonas K. A nascent polypeptide sequence modulates DnaA translation elongation in response to nutrient availability. eLife. 2021;10:e71611.

53. Rivas E. Evolutionary conservation of RNA sequence and structure. Wiley Interdiscip Rev RNA. 2021;12:e1649.

54. Vicens Q, Kieft JS. Thoughts on how to think (and talk) about RNA structure. Proc Natl Acad Sci U S A. 2022;119:e2112677119.

55. Altschul SF, Gish W, Miller W, Myers EW, Lipman DJ. Basic local alignment search tool. J Mol Biol. 1990;215:403–10.

56. Leinonen R, Sugawara H, Shumway M, International Nucleotide Sequence Database Collaboration. The sequence read archive. Nucleic Acids Res. 2011;39 Database issue:D19–21.

57. Eddy SR. Accelerated Profile HMM Searches. PLoS Comput Biol. 2011;7:e1002195.

58. Sievers F, Higgins DG. The Clustal Omega Multiple Alignment Package. Methods Mol Biol Clifton NJ. 2021;2231:3–16.

59. Edgar RC. MUSCLE: multiple sequence alignment with high accuracy and high throughput. Nucleic Acids Res. 2004;32:1792–7.

60. Waterhouse AM, Procter JB, Martin DMA, Clamp M, Barton GJ. Jalview Version 2--a multiple sequence alignment editor and analysis workbench. Bioinforma Oxf Engl. 2009;25:1189–91.

61. Guindon S, Dufayard J-F, Lefort V, Anisimova M, Hordijk W, Gascuel O. New algorithms and methods to estimate maximum-likelihood phylogenies: assessing the performance of PhyML 3.0. Syst Biol. 2010;59:307–21.

62. FigTree. http://tree.bio.ed.ac.uk/software/figtree/. Accessed 30 Jun 2022.

63. Gruber AR, Lorenz R, Bernhart SH, Neuböck R, Hofacker IL. The Vienna RNA websuite. Nucleic Acids Res. 2008;36 Web Server issue:W70-74.

64. Gasteiger E, Gattiker A, Hoogland C, Ivanyi I, Appel RD, Bairoch A. ExPASy: The proteomics server for in-depth protein knowledge and analysis. Nucleic Acids Res. 2003;31:3784–8.

65. Masquida B, Westhof E. On the wobble GoU and related pairs. RNA N Y N. 2000;6:9–15.

