## Supplemental Text for "An RNA sequence that reprograms ribosomes to bypass a 50 nucleotide coding gap is encoded by a mobile element whose sequence conservation illuminates its bypass mechanisms"

^a^ NuaBio Research, Burlingame, CA, USA.

^b^ Schools of Biochemistry and Microbiology, University College Cork, Cork T12 XF62, Ireland.

^c^ Smurfit Institute of Genetics, Trinity College, Dublin D02 VF25, Ireland

**Sequence analysis of MobA homologs**

MobA proteins are ~270-280 AA long, with an HNH endonuclease domain (~90 AA) at the N terminus, followed by a unique region with no known domains. The HNH region is 53-70% identical in protein sequence between MobA clades, and is up to 54% identical to several other T4 Mob genes, while the C-terminal region is 31-58% identical between MobA clades and has little or no detectable similarity to any other proteins. The only exception are a small set of HNH endonucleases that we designate MobZA that have extended C-terminal similarity and are typically inserted next to the dCTP pyrophosphatase gene. Members include UNI74483.1 and YP_002922087.1 (which has a NUMOD3 domain inserted, relative to other members). The C-terminus of MobE also has a short region of similarity to the C-terminus of MobA, including a strongly conserved GW.P motif.

Other Mob endonucleases have strong conservation within the HNH domain and retain a C-terminal region of similar length but with little or no sequence similarity to MobA. The HNH region is responsible for nicking the host DNA, often at a distance to the recognition site, so it may be that the C-terminal region encodes the DNA recognition site and that the sequence changes rapidly to match the appropriate recognition site. This would account for the sequence divergence between Mobs that insert at different sites, and hence have different target sequences. This theory is supported by the presence of known or predicted DNA binding domains in some other Mob proteins. For instance, *Klebsiella* phage Metamorpho has an HNH endonuclease (QPB08839) with a C-terminus homologous to the known DNA binding region of T4 I-TevI, and *Shigella* phage SSE1 (QBZ71023.1) has a C-terminal DNA binding domain related to those of GIY-YIG endonucleases. T4 MobB (NP_049674) has two predicted DNA-binding NUMOD1 domains, while T4 MobE (NP_049844.1) has a single NUMOD1 domain (as determined by its sequence similarity to both MobB domains).

**Putative VSR endonucleases in MGS11 and MGS2**

MGS11 and MGS12 have the MobA sequence of the bypass cassette replaced by a putative VSR family endonuclease. The two putative VSR proteins are 33% identical across most of their length, and have moderate similarity to many unknown and poorly-annotated sequences. Several are annotated as endonucleases, some as VSR. The largest characterized group of homologs are part of a cassette that has inserted into the DNA polymerase (gp34) of several *Aeromonas* and *Campylobacter* phage. This cassette contains intein sequences (HINT, HINTC domains) flanking the putative endonuclease, ensuring that insertion into a vital gene will allow post-translational correction of the damaged ORF by protein splicing. Other homologs of the putative VSR are also adjacent to other split genes containing intein sequences. The VSR family is very diverse in sequence, making clear identification of homologs difficult.

**Notes on sequences in File S2**

ORF1 sequences are tagged with cluster (cl) number, and NCBI accession number where available. ORF1 from Proteus phage phi-AS4 and ORF1 from Aeromonas phage CP21 have single frameshifts in the genomic sequence, which was edited to create a predicted ORF. Most ORF2 sequences are not correctly predicted from genomic annotation due to the bypass element and so do not have accession IDs.
